# The R346K Mutation in the *Mu* Variant of SARS-CoV-2 Alter the Interactions with Monoclonal Antibodies from Class 2: A Free Energy of Perturbation Study

**DOI:** 10.1101/2021.10.12.463781

**Authors:** Filip Fratev

**Affiliations:** Micar Innovation (Micar21) Ltd., Persenk 34B, 1407, Sofia, Bulgaria; Department of Pharmaceutical Sciences, School of Pharmacy, The University of Texas at El Paso, 1101 N Campbell St, El Paso, TX 79968, USA

## Abstract

The *Mu* variant of SARS-CoV-2 has been recently classified as a variant of interest (VOI) by the world health organization (WHO) but limited data are available at the moment. In particular, a special attention was given to the R346K mutation located in the receptor binding domain (RBD). In the current study we performed Free energy of perturbation (FEP) calculations to elucidate it possible impact on a set of neutralizing monoclonal antibodies (mAbs) which have been shown to be strong inhibitors of the most other known COVID-19 variants. Our results show that R346K *affects the class 2 antibodies but its effect is not so significant (0*.*66 kcal/mol); i*.*e. reduces the binding with RBD about 3 times*. An identical value was calculated also in the presence of both class 1 and class 2 antibodies (BD-812/836). Further, a similar reduction in the binding (0.4 kcal/mol) was obtained for BD-821/771 pair of mAbs. For comparison, the addition of K417N mutation, present in the newly registered *Mu* variant in July 2021 in UK, affected the class 1 mAbs by 1.29 kcal/mol reducing stronger the binding by about 10 times. Thus, the resistance effect of R346K mutation in the *Mu* variant is possible but not so significant and is due to the additional decrease of antibody neutralization based on the reduced binding of class 2 antibodies.

## Introduction

The *Mu* variant (B.1.621 lineage) of the SARS-CoV-2 (COVID-19) has been recently classified as a variant of concern (VOI) by the world health organization (WHO). It was first detected in Colombia in January 2021 and the WHO said the variant has mutations that indicate a risk of resistance to the current vaccines and stressed that further studies were needed to better understand it [1]. The *Mu* genome has a total number of 9 amino acid mutations in the virus’s spike protein, three of them located in the receptor binding domain (RBD): R346K, E484K and N501Y [2]. Whereas for the last mutations there are a plenty of data the information for the R346K substitution is limited. An initial study tested the effectiveness of sera collected from recipients of the BioNTech-Pfizer vaccine against the *Mu* variant and found that neutralization of SARS-CoV-2 B.1.621 lineage was robust, albeit at a lower level than that observed against the B.1 variant [3]. However, a recent study demonstrated that the *Mu* variant is highly resistant to sera from COVID-19 convalescents and BNT162b2-vaccinated individuals [4]. Direct comparison of different SARS-CoV-2 spike proteins revealed that *Mu* spike is more resistant to serum-mediated neutralization than all other currently recognized VOI and variants of concern (VOC). Further, it has been found that neutralizing efficacy of the BNT162b2 mRNA vaccine against the *Mu* variant has 76% neutralizing effectiveness compared to 96% with the original strain [5]. Interestingly, a two cases of *Mu* variant with an addition of the known K417N mutation was also identified in July 2021 in UK [6].

In the end of December 2020 we provided urgently high quality data about the effect of the N501Y and K417N mutations by the Free energy of binding (FEP) approach [7] and the results have been confirmed later by many experimental researches [8–10]. In the current study we used the same technique to describe the R436K mutation in the *Mu* COVID-19 variant. The FEP method is one of the most successful and precise *in silico* techniques for protein-protein interactions predictions [11]. It outperforms significantly the traditional molecular dynamics based methodologies, such as for example MM/GBSA and empirical solutions like FoldX. It also often precisely predicts the free energy differences between the mutations [12–14] and has a 90% success in the prediction whether one mutation will have either a negative or positive effect on the binding [11].

As a base of our calculations we used the recently developed SARS-CoV-2 class 1 and class 2 neutralizing monoclonal antibodies (mAbs) BD-812 (class 1), BD-836 (class 2), BD-821 (class1) and BD-771 (class 2) and also pairs of them (BD-812/BD-836 and BD-821/BD-771) for which both solid *in vitro* and structural data are available for most of the current VOCs [15]. They showed strong antiviral activity at picomol range and the obtained by cryo-EM technique molecular structures of the RBD included the E484K and K417N mutations which are both present in the last version of the *Mu* variant [6, 15].

## Results and discussion

Initially, we calculated the difference in the free energy of binding (ΔΔG) between BD-812 and the SARS-CoV-2 RBD with the R346K mutation. The result was a ΔΔG value of **0**.**66 +/-0**.**11 kcal/mol** after 5 ns of FEP/REST simulations and almost identical after 10 ns of calculations; i.e. the convergence was good (<0.3 kcal/mol/ns^-1^). The class 2 antibody BD-812 binds to the RBD with an IC_50_ of about 10 pM to all VOC [15] which can be estimated approximately to a ΔG value of about −14.9 kcal/mol (ΔG=RTln (IC_50_). Thus, our calculations predict that the R346K mutant will produce a decrease of the binding to ΔG = −14.2 kcal/mol or 33 pM which is *about 3 times reduction compared to the wild-type*. Note that by wild-type here we refer here the RBD of the *Mu* variant with all other described mutations, including K417N, but not R346K. The addition of the class 1 antibody BD-821 led to an unchanged ΔΔG value of **0.69 +/−0.12 kcal/mol** demonstrating that R346K does not have an effect on this type of mAbs.

Our simulations provide also a structural basis of the calculated reduction of the BD-812 antibody binding to the RBD. Observed conformational changes, which are located close to the Lys346 were expected and are due to the smaller size of this amino acid (**Figure 1**). The Asp53 of BD-812 switched its orientation to the RBD’s lysine 346 and consequently the Tyr27 also changed it conformation keeping almost unchanged the H-bond network of BD-812-RBD interaction surface. Thus, the hydrogen bonds and electrostatic/ionic interactions formed between Lys 346, Asp54 and Asp55 were similar to that formed by not mutated Arg residue. The main difference was that Asn450 of the mutated RBD was also involved in these interactions forming H-bonds with both Asp55 and Lys444 of the mAb by relatively small change in the conformations of these residues. The conformations of all remaining residues were almost identical (RMSD=0.6Å) in both the mutated and wild type of the RBD. Based on this data it is not unexpected that we didn’t observe a dramatic change in the free energy of binding of BD-812 to the R346K mutated SARS-CoV-2 RBD. Further, we calculated the change in the binding of the much lesser active BD-821 class 2 antibody (IC_50_ of 2-3 nM against all VOCs). The results showed a decrease in the binding by **0**.**4 +/-0**.**13 kcal/mol** *or about 2 times*. The convergence was good in this case too and the addition of class 1 antibody BD-771 to the system does not affected the result.

**Figure 1.**
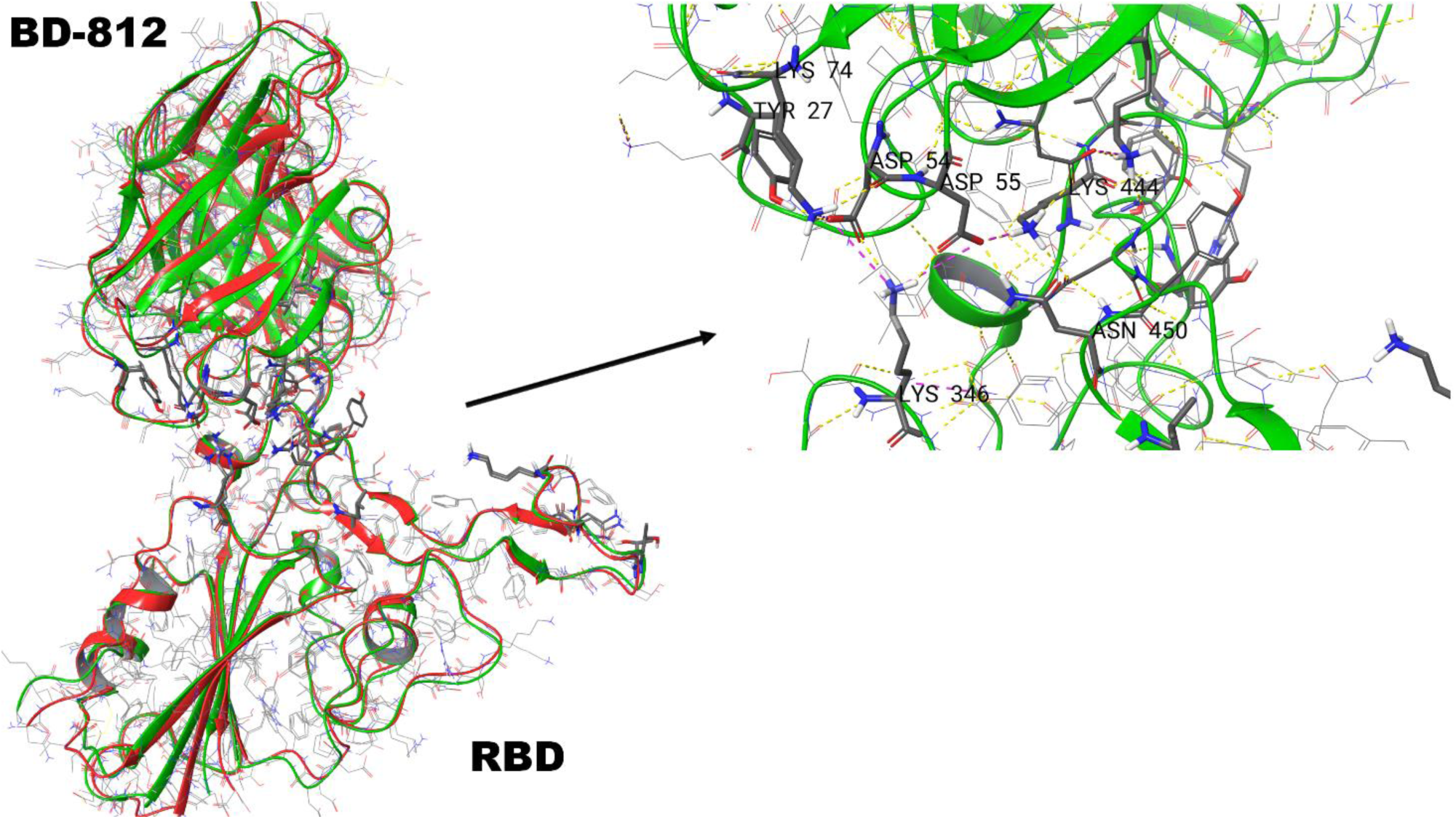
A general (*left*) and close (*righ*t) look of the identified by FEP interactions between the RBD of R346K mutant and BD-812 antibody. With green and red are shown the non-mutated and mutated RBD-BD-812 complexes, respectively. With yellow and red dot lines are shown the H-bonds and ionic interactions.

Finally, we wanted also to assay the change in the binding in the presence of K417N mutation. In December 2020 we calculated that it will have a significant impact on the class 1 antibodies [7]. This mutation was found in the Beta and also in so called Delta plus variant of the virus and has already demonstrated its significant reduction of neutralization potential for different antibodies and vaccines. Moreover, K417N has been also recently discovered in the *Mu* variant which may be a novel and potential vaccine escaping variant. The Lys417 to Asn mutation was already presented in the employed here experimental structures and we needed to make the reverse N417K mutation instead. The calculations showed that N417K increases the class 1 BD-836 antibody binding to RBD by **-1**.**29 +/-0**.**14 kcal/mol** *which is about 10 times compared to the original Mu variant*. This charged residue is located right on the class 1 antibodies binding surface. The simple substitution of Asn with Lys in the used experimental structure lead to the conclusion that Lys417 stabilizes the interactions with Ser30 and Glu74 of the antibody. However, the REST MD simulations demonstrated that the Ser30-Glu74 contact is not so stable and Glu74 moves toward Arg72 (**Figure 2**). This movement is restricted by the introduction of Lysine and as a result the Glu74 (also the all loop) is closer to RBD and formed a strong Lys417-Glu74 H-bond. The better BD-836-RBD contact and this hydrogen bond lead to the observed increase of the binding. Thus, the significant impact of this mutation detected in UK [6] was confirmed by the new calculations and we can conclude that the action of such COVID-19 *Mu* variant will be more severe and should be closely monitored.

**Figure 2.**
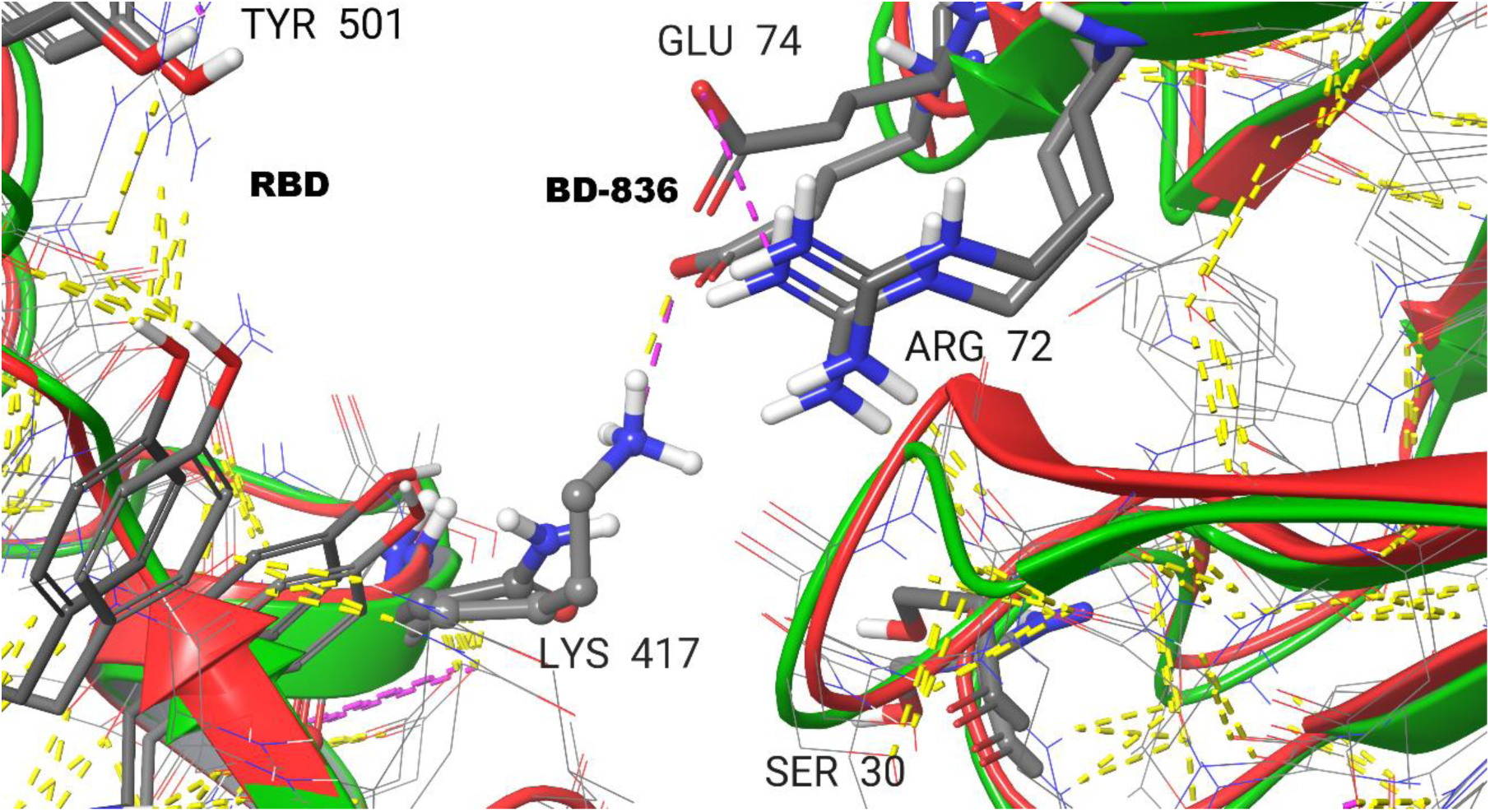
Identified interactions between RBD and class 1 antibody BD-836. With red and green are shown the non-mutated and mutated RBD-BD-836 complexes, respectively. With yellow and red dot lines are shown the H-bonds and ionic interactions.

As a limitation of this study one can argue that we used a small number of antibodies and eventually the R346K mutation can act a bit different in other cases. However, a drastic difference is unlikely to be observed.

## Methods

The methods used have been described in details previously [7]. In short, to perform our FEP calculations and structural studies we used the recently deposited structures with pdb accession numbers 7EZV (for the RBD-BD-812/836 complex) and 7EY5 (for the RBD-BD-821/836 complex), respectively. For the FEP assessments of the K417N mutation we used the pdb id 7EZV structure too. All protein preparations and calculations were performed by Schrödinger suite software [16]. To calculate the differences in the free energy of binding for each complex in this study we employed the Desmond FEP/REST approach described in details previously [11–14]. A sample scheme of the thermodynamical cycle for the calculations of the binding affinity change due to mutations in interacting protein-protein interface is shown on Figure 1 in ref [17]. The binding free energy change ΔΔG_AB_ or in simple ΔΔG can then be obtained from the difference between the free energy changes caused by the particular mutation in the bound state (ΔG_1_, complex leg) and the unbound state i.e. in solvent (ΔG_2_, solvent leg). In a typical FEP calculation for a mutation from state A to state B, several perturbation lambda (λ) windows are needed in order to obtain a smooth transition from the initial state A to the final state B. The default sampling FEP+ protocol was applied here with the number of λ windows either 12 or 24, in dependence of the mutation charge; i.e. same or different. An equilibration and 5 ns-long replica exchange solute tempering (REST) simulations in a muVT ensemble was further conducted. Only the mutated atoms was included in the REST “hot region” region. OPLS4 force field was used for the all simulations [18]. The obtained average structures from the simulations were used for the structural comparison on **Figures 1** and **2**. All results can be reproduced and further analyzed by the Appending I files, which include the structures, all parameters and all protocols, which are available on request.

## Acknowledgments

Thanks due to the Suman Sirimulla for the helpful discussion.

